# Genomic loci for sclerotinia stem rot resistance and chlorophyll stability in *Brassica napus*: integrating GWAS with microbiome insights

**DOI:** 10.1101/2024.08.20.608782

**Authors:** Aakash Chawade, Vishnukiran Thuraga, Siim Samuel Sepp, Samrat Ghosh, Farideh Ghadamgahi, Firuz Odilbekov, Saraladevi Muthusamy, Ramesh R Vetukuri, Kibrom B. Abreha

## Abstract

Sclerotinia Stem Rot (SSR) disease is one of the most serious diseases affecting the yield and quality of oilseed rape (*Brassica napus*). Understanding the genetic basis of the resistance trait in oilseed rape to SSR and microbiome composition for enhanced resistance is crucial for developing resistant varieties and sustainably mitigate the impact of the disease. In this study, in a panel of 168 oilseed rape accessions, most resistant (NGB 13503 and NGB 13834) and susceptible (NGB 13497 and NGB 13897) accessions are identified. A Genome-wide association study (GWAS) identified 47 SNPs linked to the SSR lesion length, lesion area, and lesion relative to the leaf area. Among the SNPs significantly linked to lesion length were Bn-A04-p10555408, Bn-A07-p12487549, Bn-A09-p4652268, Bn-A09-p4916858 and to our knowledge, these SNPs have not been previously linked to SSR resistance in oilseed rape. Moreover, the study identified 24 SNPs linked with chlorophyll content before SSR inoculation (SPADH), after the SSR inoculation (SPADI), and chlorophyll index (CI). Maintaining the chlorophyll level is correlated with the SSR resistance. Furthermore, bacterial taxa (e.g. *Pseudomonas*, *Methylobacterium*, and *Aquabacterium*) and fungal taxa (e.g. *Mycosphaerellales*, *Thelebolales*, and *Akanthomyces*) were enriched in the resistant compared to in the susceptible oilseed rape accessions. The SNPs linked to lesion length showed consistent haplotype variation between these selected accessions. Given the absence of complete resistance against SSR, the study provides insights into the significance of maintaining chlorophyll levels and considering microbiome composition for enhancing the level of existing partial resistance to SSR in oilseed rape.

## Introduction

Oilseed rape (*Brassica napus* L.), formed by natural interspecies hybridization of *Brassica rapa* (AA; 2n = 20) and *B. oleracea* (CC; 2n = 18), is an economically important allotetraploid (AACC; 2n = 4x = 38) oilseed crop. Currently, oilseed rape is cultivated globally providing an estimated annual production exceeding 75 million tonnes (FAOSTAT, 2023). It is one of the most commonly used vegetative oils along with soybeans and palm oil. The crop has high seed protein content and high seed oil yield and quality, which is rich in mono and polyunsaturated fatty acids and tocopherols (Stepien et al., 2017; Rout et al., 2018; Chew, 2020). As a result, oilseed rape is used as food, a protein-rich feed for animals, and has multiple industrial applications (Raboanatahiry et al., 2021).

The small number of founder parents involved in allopolyploidization events in oilseed rape, strong breeding selection for high seed oil quality, and intensified cultivation for higher yield have limited the oilseed rape genetic pool (Wu et al., 2014; Liu et al., 2022). This has contributed to various fungal diseases causing serious losses in oilseed rape yield and quality (Van de Wouw et al., 2016). Particularly, the fungal pathogen *Sclerotinia sclerotiorum*, the causal agent of the devastating Sclerotinia Stem Rot (SSR) disease, is present in all major oilseed rape growing areas and causing significant yield and quality losses in oilseed rape (Derbyshire and Denton-Giles, 2016; Neik et al., 2017). SSR has a broad host range including several Brassica oilseed crops (Ficke et al., 2018), indicating its significant importance in oilseed production in general. Up to 80% yield loss due to SSR disease has been reported in major oilseed rape growing areas (Wu et al., 2016). The emergence of virulent and fungicide resistant strains of the pathogen could lead to frequent outbreaks in oilseed rape growing areas (Derbyshire and Denton-Giles, 2016). Therefore, there is a pressing need to develop effective and sustainable methods to reduce SSR disease pressure, and consequently minimize seed yield and quality losses.

Enhancing the oilseed rape resistance against SSR disease through genetic improvement is the most environmentally sustainable approach for managing SSR. Identifying resistance genotypes with complete resistance against SSR in diverse oilseed rape germplasm is essential for developing resistant varieties. Despite considerable breeding efforts, however, oilseed rape varieties with complete resistance against SSR have not been developed so far (Neik et al., 2017; Ding et al., 2021). This is partly attributed to the aggressiveness of the pathogen employing diverse strategies to overcome the host resistance, and limited understanding of the oilseed rape-SSR pathosystem, and genetic regulation of the host resistance.

Multigenic resistance governed by quantitative trait loci (QTLs) is the most important form of oilseed rape resistance against SSR (Wu et al., 2016; Neik et al., 2017). Nevertheless, only a few QTLs related to SSR resistance have been identified so far, mainly using bi-parental mapping populations of the crop (Derbyshire and Denton-Giles, 2016; Wu et al., 2016). Bi-parental population-based QTL mapping is only restricted to the diversity and recombination events of between a few founder genotypes thus resulting in low resolution of the genetic map (Dahanayaka and Martin, 2023). Consequently, the QTLs identified so far have limited application for improving the SSR resistance in oilseed rape.

Genome-wide association study (GWAS), which takes advantage of the phenotypic variation and historical recombination in natural populations, as a powerful tool to identify quantitative trait loci (QTLs) linked to important traits of interest (Korte and Farlow, 2013). GWAS uses thousands of SNPs generated through next-generation sequencing technologies and it is instrumental for establishing marker-trait association (MTA) and the development and validation of genetic markers for use in Markers assisted selection (MAS). Indeed GWAS has been successfully used to identify QTLs linked to disease resistance (Wu et al., 2016; Roy et al., 2021), as well as other important agronomic traits such as yield and oil quality-related traits in oilseed rape (Zhao et al., 2022; Xiang et al., 2023).

Host resistance is a result of the host, pathogen, and environment interactions involving biotic and abiotic factors. Pathogen infection significantly reduces the photosynthetic performance and subsequently the yield and quality. Maintaining the photosynthetic capacity could be one mechanism of SSR resistance in oilseed rape, as recently shown possible involvement of photosynthesis in silicon (Si) induced resistance against the pathogen (Feng et al., 2021). Chlorophyll levels could be used as a proximal trait to measure disease in crops (Odilbekov et al., 2018; Koc et al., 2022). Moreover, with the advent of sequencing technologies and analysis tools, there is growing evidence regarding the role of microbes affecting host-pathogen interactions (Rybakova et al., 2017; Liu et al., 2021). Since no complete resistance against SSR has been identified in diverse germplasm screening or developed through breeding, understanding the genetic basis for maintaining the photosynthesis level and microbiome composition would be invaluable for enhancing the resistance and developing effective disease management strategies.

In the present study, we have identified oilseed rape genotypes with partial resistance against SSR, and identified marker-trait associations (MTAs) for the SSR-resistance and chlorophyll levels during the SSR infection. Furthermore, using the most relatively resistant and susceptible genotypes, we showed differential microbiome composition in these genotypes and established the correlation to the SSR resistance MTAs identified using GWAS.

## Materials and methods

### Plant material and experimental setup

A panel of 168 oilseed rape accessions, obtained from the Nordic Genetic Resource Centre (NordGen), consisting mostly of varieties from Nordic origin was used for Genome-wide analysis study (GWAS) analysis to identify genomic regions linked to SRR resistance (Supplementary Table S1). Ten plants from each accession were grown in seedling trays and at three weeks, leaves from every plant were collected from each accession for Genotyping-by-sequencing (details below). Three seedlings per accession were each transferred into 2L potting soil separately and arranged in a completely randomized (CRD) design with three replications in a controlled climate chamber facility with adjustable climate conditions at the Swedish University of Agricultural Sciences at Alnarp. The plants were grown in the chamber under long-day conditions (16/8h photoperiod, 25/20°C Day/night, and 65% RH) until evaluated for resistance against SSR using detached leaf assays.

### Fungal strain maintenance and SSR resistance evaluation

*S. sclerotinia* isolate was obtained from Lantmännen and aseptically maintained on potato dextrose agar (PDA) at 4°C in the dark. A five mm plug was placed in the middle of freshly prepared PDA plates and incubated at room temperature in the dark. Agar plugs (3mm Ø) were punched from the actively growing edge of three-day-old fungal PDA plates. For detached leaf assays, three leaves per accession per plant were excised from the 6-week-old plants, placed in a transparent plastic box (39 x 29 x 12.5 cm), and humidified to 100% with a water-soaked gauge to maintain higher humidity. The gauge was separated from the leaves by a plastic net. One fungal plug (3mm Ø) on both sides of the midrib on abaxial surface of the detached leaves. Boxes with inoculated detached leaves were sealed with a parafilm, incubated at room temperature and 16:8 photoperiod. Lesion diameter and leaf pictures were taken at 3 days post inoculation (3 dpi).

### Phenotyping and data analysis

The diameter of the lesion area (Lesion Length, LL) on the detached leaves was measured manually at 3 days post inoculation (dpi). Immediately, pictures of each leaf were taken using a mounted NIKON D3500 (Nikon, Tokyo), and the pictures were analyzed on a leaf basis to measure the lesion area (LA) and calculate the relative lesion area (RLA) using ImageJ. According to this measurement, the ten most relatively resistant and ten most susceptible accessions to SSR were selected for microbiome recruitment analysis (Details below).

Leaf chlorophyll concentration before infection (SPADH) was measured immediately after the leaves were detached and placed in the transparent infection boxes. Leaf chlorophyll concentration was also measured at 3 dpi after infection (SPADI). Both SPADH and SPADI were measured with the MC-100 chlorophyll meter (Apogee Instruments Inc., Logan) at five locations of each leaf and average data was expressed as SPAD values. The chlorophyll index (CI) was calculated in terms of chlorophyll loss under infection conditions by employing the following formulae.

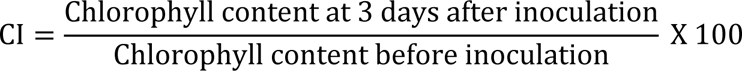

### SNP array genotyping of oilseed rape accessions

One leaf from each of the 10 plants from each accession was collected, piled, and punched with 3mm cork. The 3mm discs of each accession were placed in 96 well plates provided as part of the BioArk^TM^ Leaf collection kit from LGC Biosearch Technologies (Berlin, Germany). During sampling, the well plates were placed on ice to slow down the physiological process and drying of excised discs. Upon collecting the leaves from all accessions, the plates were covered with cotton incubated in a freeze dryer for three days and sent to the sequencing company. The genotyping was done using the Brassica 19K Illumina SNP array (Illumina Inc, San Diego CA). A total of 18579 informative single nucleotide polymorphism (SNP) markers, representing both the *B. rapa* A-genome and *B. oleracea* C-genome, were obtained. Of which, 14 266 SNPs were mapped to the *B. napus* reference genome (reference) and marker distribution across the genome was visualized using the Scientific and Research plot tool (SRPlot) (Tang et al., 2023). A total of 12 738 high-quality (missing data < 10%, MAF > 0.05) were filtered in TASSEL v.5.2.86 (Bradbury et al., 2007), and used for further analysis.

### Linkage disequilibrium (LD) and population structure

Using the 12 738 high-quality SNPs, the pairwise LD values (*r*^2^), which refer to the nonrandom association of alleles at different loci, were calculated (window size = 50 markers) in TASSEL v.5.2.86 (Bradbury et al., 2007). The LD (*r*^2^) values were plotted against the genetic distance (bp) between marker pairs for each chromosome using the LD score plotting script in R Studio as described previously (Remington et al., 2001). The pattern of genome-wide LD decay was considered at *r*^2^= 0.1 threshold, which is at half compared to the maximum LD (*r*^2^) value.

### Genome-wide association study (GWAS) analysis

Best linear unbiased predictions (BLUPs) for the measured traits from the 168 oilseed rape accessions were estimated using the Multi Environment Trial Analysis with R (META-R) (Alvarado et al., 2020), and used as a phenotype and their corresponding genotyping data was used for the GWAS analysis. The GWAS analysis was performed with the most robust models bayesian-information and linkage-disequilibrium iteratively nested keyway (BLINK) (Huang et al., 2018) and fixed and random model circulating probability unification (FarmCPU) (Liu et al., 2016) with GAPIT Version 3 in RStudio (Wang and Zhang, 2021). The relative kinship matrix (K) was computed with the VanRanden method (VanRaden, 2008), population structure was visualized with a heatmap performed in GAPIT in R (Wang and Zhang, 2021). The suitability of the models BLUPs for the GWAS analysis was assessed using histograms, quantile-quantile (Q-Q) plots, and scatter plots. SNPs with *p* < 0.001 [–log_10_(*p*) > 3] were considered significantly associated with the phenotypic traits measured.

### Effect of the significant SNPs on the phenotypes measured

All the oilseed rape accessions were grouped according to the allele variation at the most significant SNPs associated with the traits measured. To determine the significance among groups formed based on favorable and alternative allele, a two-sample t-test was conducted in R.

### Microbiome recruitment in relatively resistant and susceptible accessions of oilseed rape DNA extraction, amplification and sequencing

DNA was extracted from the collected leaf samples of the most relatively resistant and susceptible accessions using a modified DNeasy PowerSoil Pro kit (Qiagen, CA, USA) protocol, as mentioned below, with three replications per accession. The frozen leaf samples were ground to the powder in a pre-chilled mortar and pestle filled with liquid nitrogen before proceeding with the manufacturer’s recommended protocol. The wash step with solution EA was repeated three times to ensure the removal of phenolic compounds. The extracted DNA’s quality and quantity were evaluated using a Nanodrop (Nanodrop 8000 Thermo Fisher Scientific, USA).

For bacteria, the V5-V6 regions of the bacterial 16S rRNA gene were amplified using the primer pair 799F (AACMGGATTAGATACCCKG) and 1115R (AGGGTTGCGCTCGTTRC). For fungi, the internal transcribed spacer 1 (ITS1) region was targeted by the primer pair ITS1Kyo2F (TAGAGGAAGTAAAAGTCGTAA) and ITS86R (TTCAAAGATTCGATGATTCA). For each sample, PCR reactions were performed using a 20 μL mixture comprising 1x MyTaq buffer containing 1.5 units of MyTaq DNA polymerase (Bioline GmbH, Luckenwalde, Germany), two μL of BioStabII PCR Enhancer (Sigma‒Aldrich Co.), ∼1-10 ng of template DNA and 15 pmol of the respective forward and reverse primers, as mentioned above. PCR was carried out using the following program: predenaturation at 96°C for 1 minute, denaturation at 96°C for 15 seconds, annealing at 55°C for 30 seconds and extension at 70°C for 90 seconds, followed by 30-33 cycles of amplification of prokaryotic genes and 35-40 cycles of amplification of eukaryotic genes. The amplified PCR products were purified with Agencourt AMPure beads (Beckman Coulter, Brea, CA, United States). Illumina libraries were constructed using the Ovation Rapid DR Multiplex System 1-96 (NuGEN Technologies, Inc., California, USA) with 100 ng of purified amplicon pool DNA for each type. After library construction, the Illumina libraries (Illumina, Inc., CA, USA) were combined and subjected to size selection through preparative gel electrophoresis. Sequencing was performed on an Illumina MiSeq platform at the LGC’s sequencing facility in Berlin (LGC Genomics GmbH, Germany).

### Amplicon data processing and analysis

After sequencing, primers, and adapters were removed from amplicon data using cutadapt v4.7 (Walker et al., 2014), the quality of the reads was improved with the same tool. Next, high-quality filtered amplicon reads were demultiplexed with QIIME2 v2022.8 (Bolyen et al., 2019), and the DADA2 plugin (Callahan et al., 2016) of QIIME2 was employed to denoise, dereplicate and remove chimeric reads. The resulting amplicon sequence variants (ASVs) were taxonomically classified using the pre-trained 16S rRNA database SILVA v138.1(Quast et al., 2013) and the ITS database UNITE v9 (Nilsson et al., 2019). A Naive Bayes classifier was used to train databases. ASVs were unassigned, and ASVs were assigned to chloroplast, and mitochondria were removed.

### Statistical analysis

R v4.2.0 (Team R Core, 2013) was used for all the statistical analysis. A phyloseq object comprising ASVs table, taxonomy, sequence and sample metadata file was made using the R package “phyloseq v1.26.1” (McMurdie and Holmes, 2013). Before statistical analysis, phyloseq object was rarefied. The plot_taxa_composition () function of the “microbiome v3.19”(Leo Lahti, 2017) and “microbiomeutilities v0.99” (Lahti, 2018) packages were used to plot the relative abundance (%) of taxons. Data normality was checked with Shapiro.test () function from the “stats v3.6.2” (Team, 1970) package. Alpha diversity metrics “Chao1” and “Shannon” were computed with “vegan v2.5.1” (Dixon, 2003), and the same package Wilcoxon rank sum test was performed to check the difference in the sample groups. Linear discriminatory analysis (LDA) was performed with “microbiomeMarker v1.2.1” package (Cao et al., 2022), cut-off value for LDA was set at 2 and *p-*value at <0.05.

### Data availability

All the amplicon data (16S&ITS) were submitted into the sequence read archive repository under BioProject: Id PRJNA1063738.

## Results

### Phenotypic variability within the oilseed GWAS panel

A GWAS panel composed of 168 diverse oilseed rape accessions was evaluated for Sclerotinia stem rot (SSR) lesion length (LL), lesion area (LA), and relative lesion area (RLA). Leaf chlorophyll concentration was measured before inoculation (SPADH) and after inoculation (SPADI) with SSR respectively, and the chlorophyll index (CI) in these accessions was calculated using these measurements. The summary statistics such as mean, range, and coefficient of variance for the traits measured are presented in Table 1.

**Table 1.** Mean and coefficient of variation for the measured traits.

| Trait measured | Mean ± SD | Range | CV (%) |
| --- | --- | --- | --- |
| Lesion length (LL) (mm) | $58.45 \pm 1.56$ | 53.53 - 63.87 | 29 |
| Lesion area (LA) (cm<sup>2</sup>) | $19.46 \pm 0.55$ | 17.66 - 22.61 | 56 |
| Relative lesion area (RLA) (%) | $48 \pm 0.02$ | 45 - 57 | 52 |
| Chlorophyll (Healthy) - SPADH | $37.21 \pm 0.94$ | 34.63 - 40.11 | 9 |
| Chlorophyll (Inoculated) - SPADI | $18.86 \pm 0.91$ | 15.25 - 22.06 | 24 |
| Chlorophyll index (CI) (%) | $50.36 \pm 1.98$ | 41.47 - 56.65 | 21 |

The mean LL was 58.5 ± 1.56 mm with the shortest 53.5 mm and the longest 63.9 mm whereas LA ranges from 17.7 - 22.6 cm^2^ with a mean value of 19.5 ± 0.6 cm^2^. On average, the lesion covered 48% of the inoculated leaf area (RLA). The CV was higher for LA (56%) and RLA (52%) compared to LL (29%), highlighting the importance of measuring various aspects of the lesion growth and leaf to understand the growth of the disease on inoculated leaves. We found a strong positive correlation (r^2^ = 0.60) between LL and LA (Figure 1), however, RLA showed a weak correlation to these SSR resistance traits (Figure 1).

**Figure 1.**
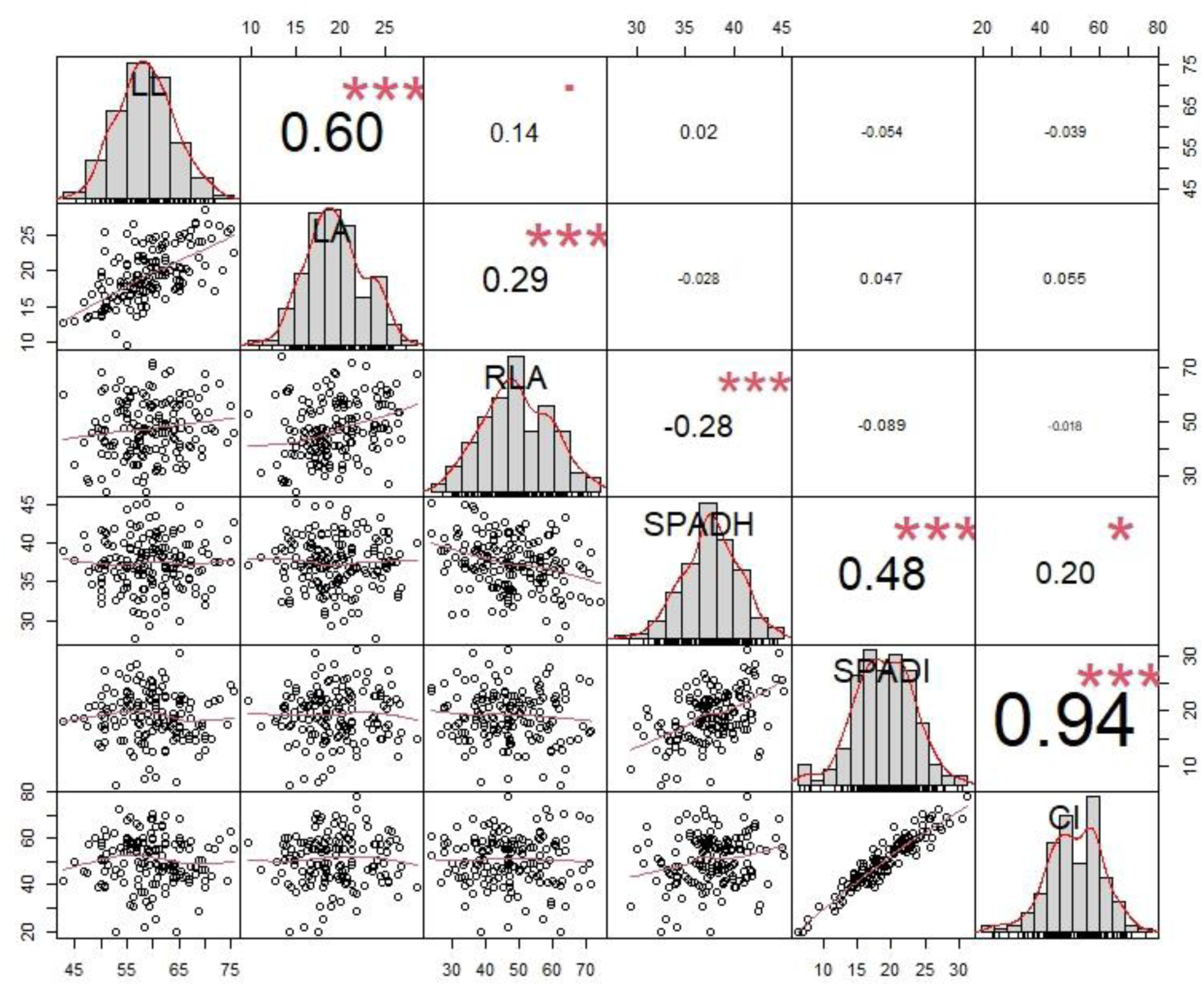
Frequency distribution, bivariate scatter plots, and correlation analysis among the traits measured in 168 accessions of oilseed rape. Sclerotinia stem rot (SSR) lesion length (LL), lesion area (LA), and the relative leaf area (RLA) measured at 3 days post inoculation (dpi). Leaf chlorophyll concentration was measured before inoculation (SPADH) and at 3 dpi (SPADI) with SSR respectively, and the chlorophyll index (CI) was calculated from both measurements. Significance level of the correlation values: *** (*P* < 0.001), **(*P* < 01), *(*P* <0.05),. (*P* < 0.1).

Although there was no correlation (Figure 2), the RLA reflects the dropping of the chlorophyll content due to the infection from 37.21 in SPADH to 18.86 in SPADI. CI value ranges from 41.47-56.65% (Table 1) and it is significantly correlated to SPADI (r^2^ = 0.94) while SPADH has little effect (r^2^ = 0.20) (Figure 1). The CV for SPAD increased from 9% in SPADH to 24% in SPADI indicating the chlorophyll content in the accessions was affected differently by the SSR inoculation.

**Figure 2:**
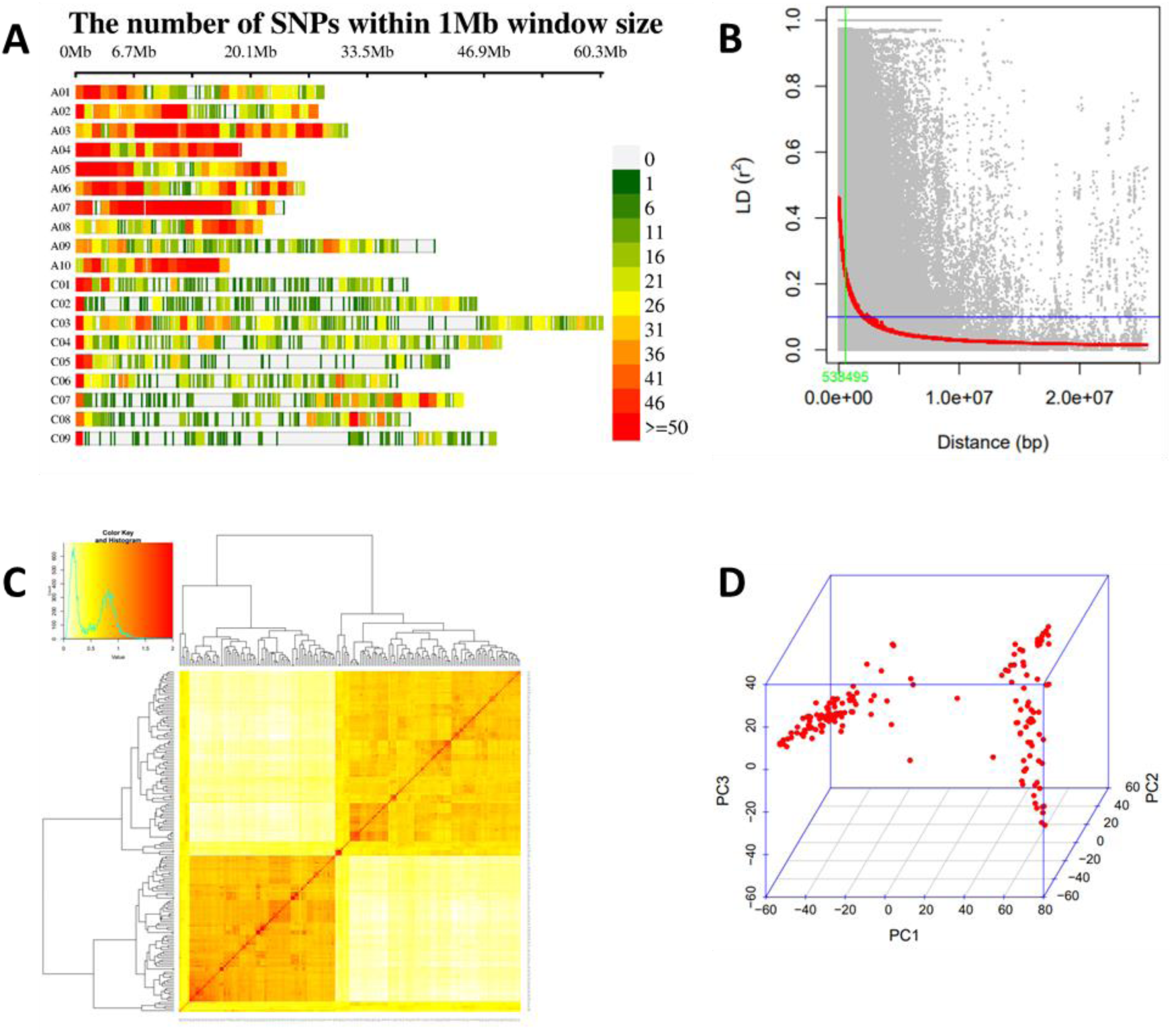
(A) Number of SNPs within 1Mb window size of *Brassica napus* genome; B) PCA; *(C) Kinship plot as a heatmap of the VanRaden kinship matrix*; (D) *LD decay scatter plot with r^2^ values of pairwise SNPs*.

### SNPs distribution, LD decay, and population structure

The high-quality 12 738 SNPs distributed throughout the A-genome (7713) and C-genome (5025) were visualized with SRPlot. A non-uniform pattern of SNPs distribution per 1 million base window was observed across the 19 chromosomes with the most SNPs mapped on chromosome A03 (1282) and least on chromosome C01 (536).

The LD squared correlation coefficient (*r^2^*), for all the 1 237 high-quality SNP markers, was used for estimating the extent of LD decay. Genome-wide LD decayed to *r*^2^ = half decay at 0.53 Mb (Figure 2B). The LD decay varied between 0.39Mb for A- and 1,07Mb for C-subgenomes as well as across the chromosomes with the longest 4,83Mb for C01 and the shortest 0.25Mb for A07 (Table 2). Based on this LD decay, all significant SNPs within LD decay distance values in each chromosome were considered as part of the same QTL.

**Table 2.**
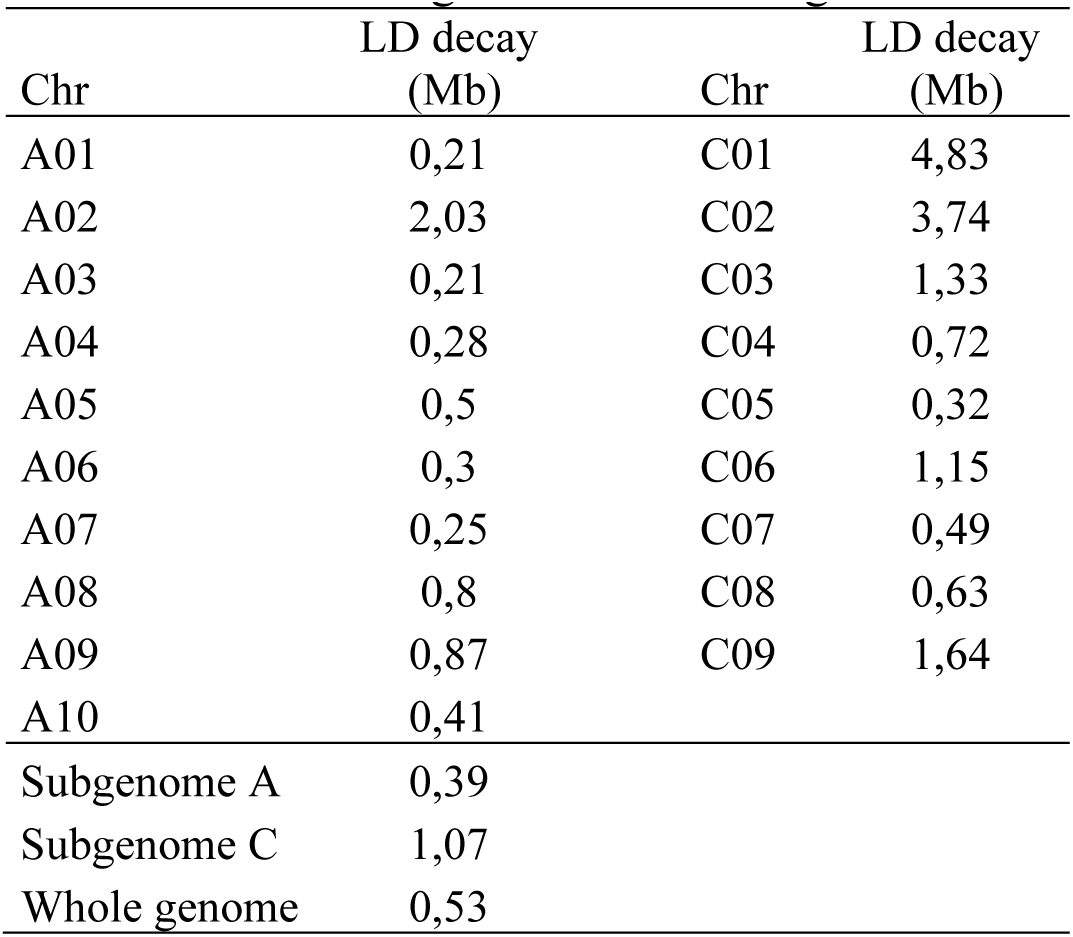
Linkage disequilibrium for the chromosomes in subgenome A and subgenome C.

| Chr | LD decay (Mb) | Chr | LD decay (Mb) |
| --- | --- | --- | --- |
| A01 | 0,21 | C01 | 4,83 |
| A02 | 2,03 | C02 | 3,74 |
| A03 | 0,21 | C03 | 1,33 |
| A04 | 0,28 | C04 | 0,72 |
| A05 | 0,5 | C05 | 0,32 |
| A06 | 0,3 | C06 | 1,15 |
| A07 | 0,25 | C07 | 0,49 |
| A08 | 0,8 | C08 | 0,63 |
| A09 | 0,87 | C09 | 1,64 |
| A10 | 0,41 |  |  |
| Subgenome A | 0,39 |  |  |
| Subgenome C | 1,07 |  |  |
| Whole genome | 0,53 |  |  |

A clustering heat map created using a kinship matrix and Principal components analysis structured the 168 accessions into two subpopulations (Figure 2C and 2D). In both analyses, the accessions were mostly grouped according to their growth habit, winter accessions or spring accessions. PC1 and PC2 explain 34.5% and 4% of the total variation, respectively (Figure 2D). The grouping in principal components analysis corresponds with the VanRaden kinship matrix clustering.

### Genome-wide marker-trait associations

The main objective of this study was to identify SNPs significantly related to the seed, chlorophyll, and SSR resistance traits in the oilseed rape accessions. The normal distribution of the histograms (Figure 1) indicated the suitability of the traits’ BLUPs for GWAS analysis. For the six traits measured, a total of 71 significant MTAs, 34 in A and 37 in C sub-genomes, were detected with the FarmCPU and BLINK models (*p*-value < 0.001). Except for ten MTAs associated with the chlorophyll index, all the MTAs associated with the other traits were detected in both GAPIT models. The MTAs were distributed across the ten chromosomes of the A subgenome and seven chromosomes of the C subgenome, except for chromosomes C01 and C09. The highest number of MTAs were located on Chr-C02 (17MTAs) followed by chromosome A01 (11 MTAs). The MTAs detected on C02 were associated with the chlorophyll index, chlorophyll after infection (SPADI), and lesion length. Some MTAs chromosome A01 and C02 were also associated with relative lesion area. Except for LA, all the traits had MTAs from subgenomes A and C.

### MTAs for Sclerotinia Stem Rot (SSR) resistance traits in oilseed rape

A total of 47 MTAs on 12 chromosomes, six from each sub-genome, were found significantly (p < 0.001;–log10(p) > 3) associated with the Sclerotinia stem rot (SSR) resistance traits lesion length (LL), lesion area (LA) and the relative lesion area (RLA) (Table 3 and Figure 3). Among the ten MTAs linked with the LL were two on chromosome C04 and two on C08 as well as two MTAs, Bn-A09-p4652268 and Bn-A09-p4916858 on chromosome A09.

**Figure 3.**
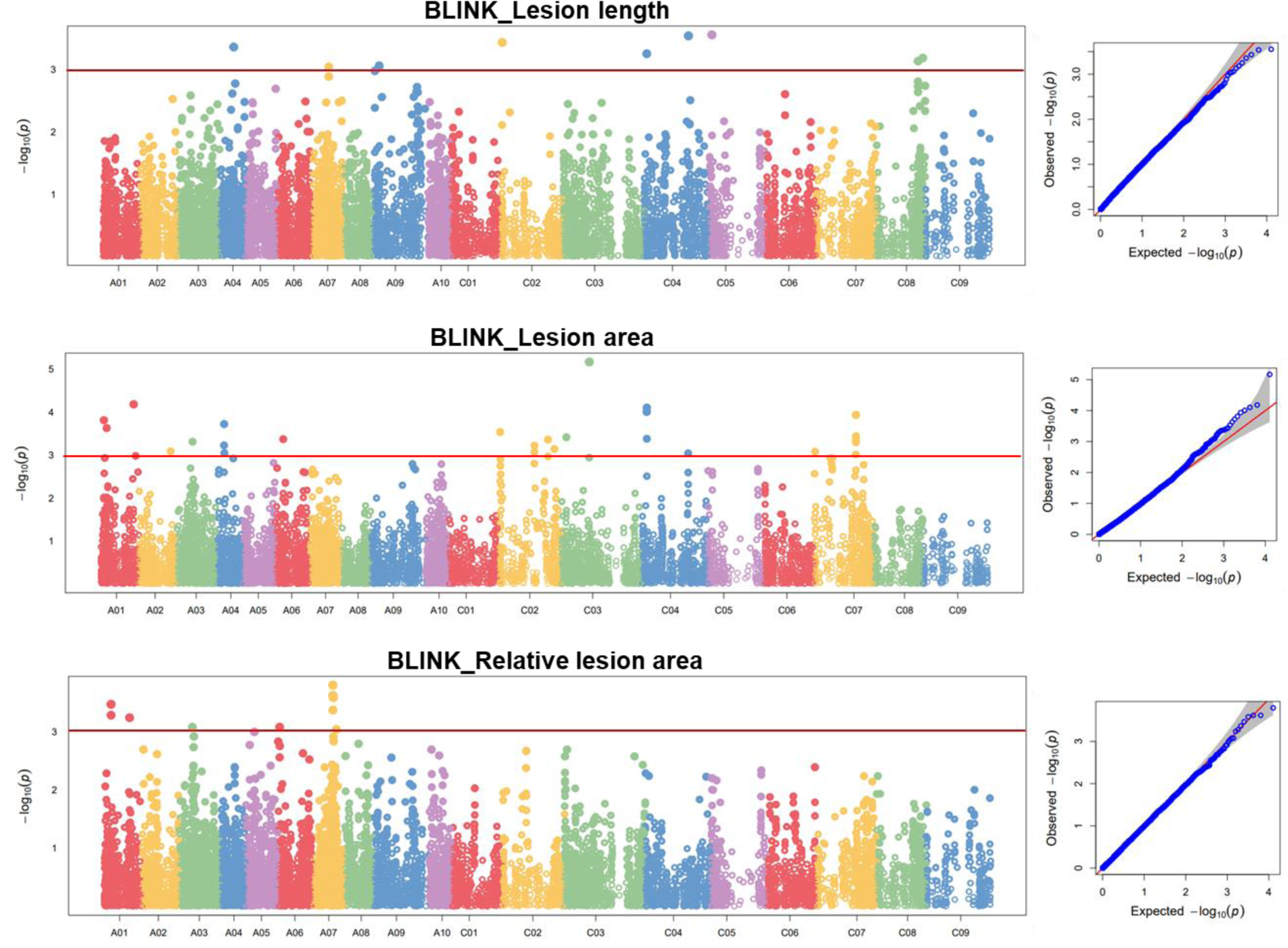
Manhattan (left) and Q-Q (right) plots for the GWAS analysis of sclerotinia stem rot (SSR) resistance-related traits Lesion length, Lesion area, and relative lesion area in oilseed rape accessions.

For the LA, nine MTAs were found in the A sub-genome across five chromosomes (A01, A02, A03, A04, and A06) and 17 MTAs were in the C sub-genome distributed across four chromosomes (C02, C03, C04, and C07). Based on an average LD decay of 0.53 Mb, thirteen MTAs linked to the SSR-IA were classified into four QTLs with three MTAs on chromosome A04 (48.49 - 51.37 Mb), two MTAs on C02 (25.69 – 25.72 Mb), three MTAs on C04 (32.45 – 32.85Mb), and five MTAs on C07 (31 – 31-13Mb) (Table 3). Other MTAs associated with LA were Bn-A01-p2291940 (2. 29 Mb), Bn-A01-p24575577 (24.58 Mb), and Bn-A01-p4353657 (4. 35 Mb) on chromosome A01 (Table 5). All the MTAs associated with the RLA were located in the A sub-genome. Among the three MTAs were Bn-A01-p5265598 (5.26 Mb) and Bn-A01-p5269368 (5.26 Mb) (Table 3). Similarly, four MTAs found at 14.76 Mb (Bn-A07-p14766907 and Bn-A07-p14768265), 14.77 Mb (Bn-A07-p14770619), and 14.79 Mb (Bn-A07-p14792267) could be considered one QTL.

**Table 3.**
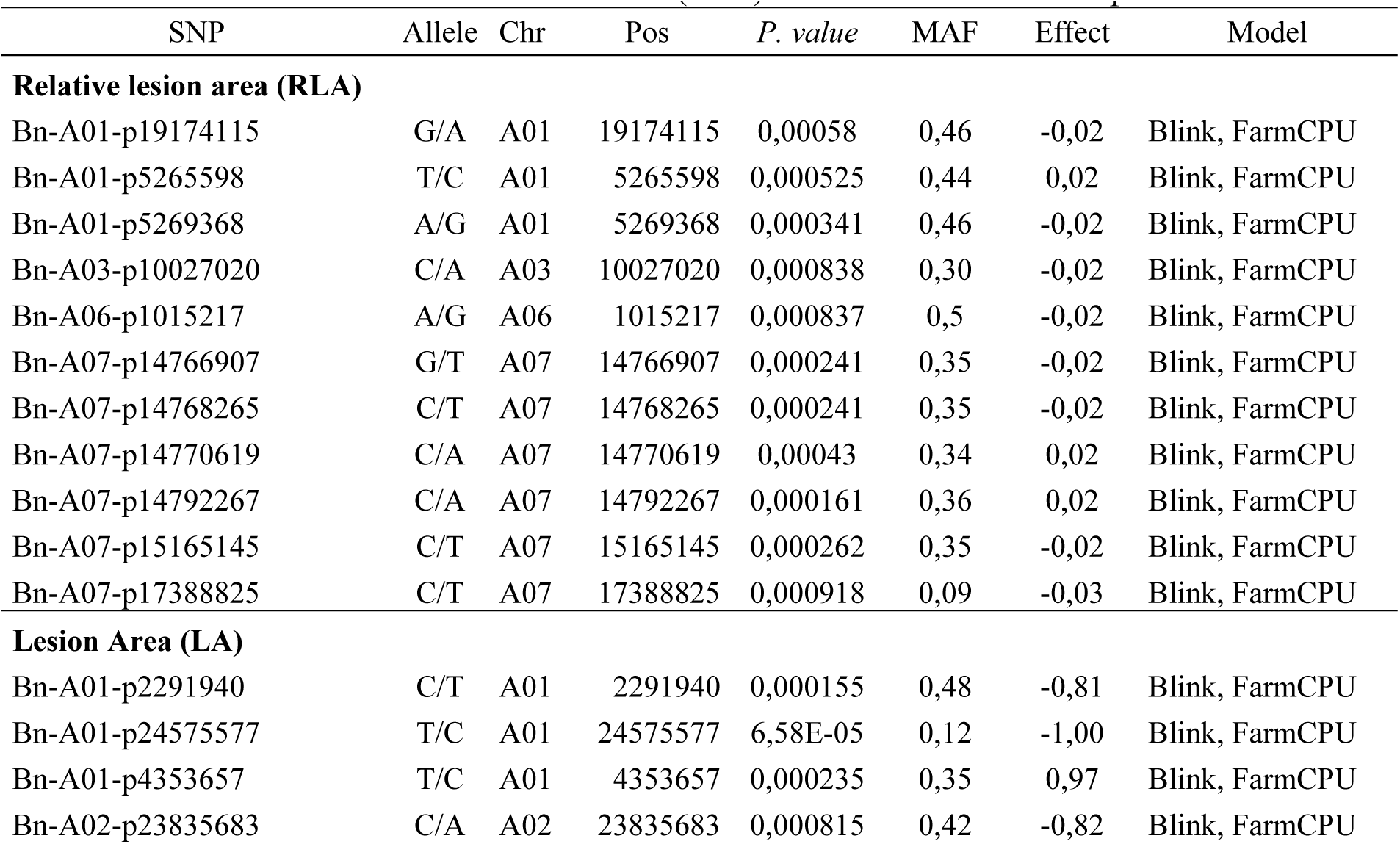
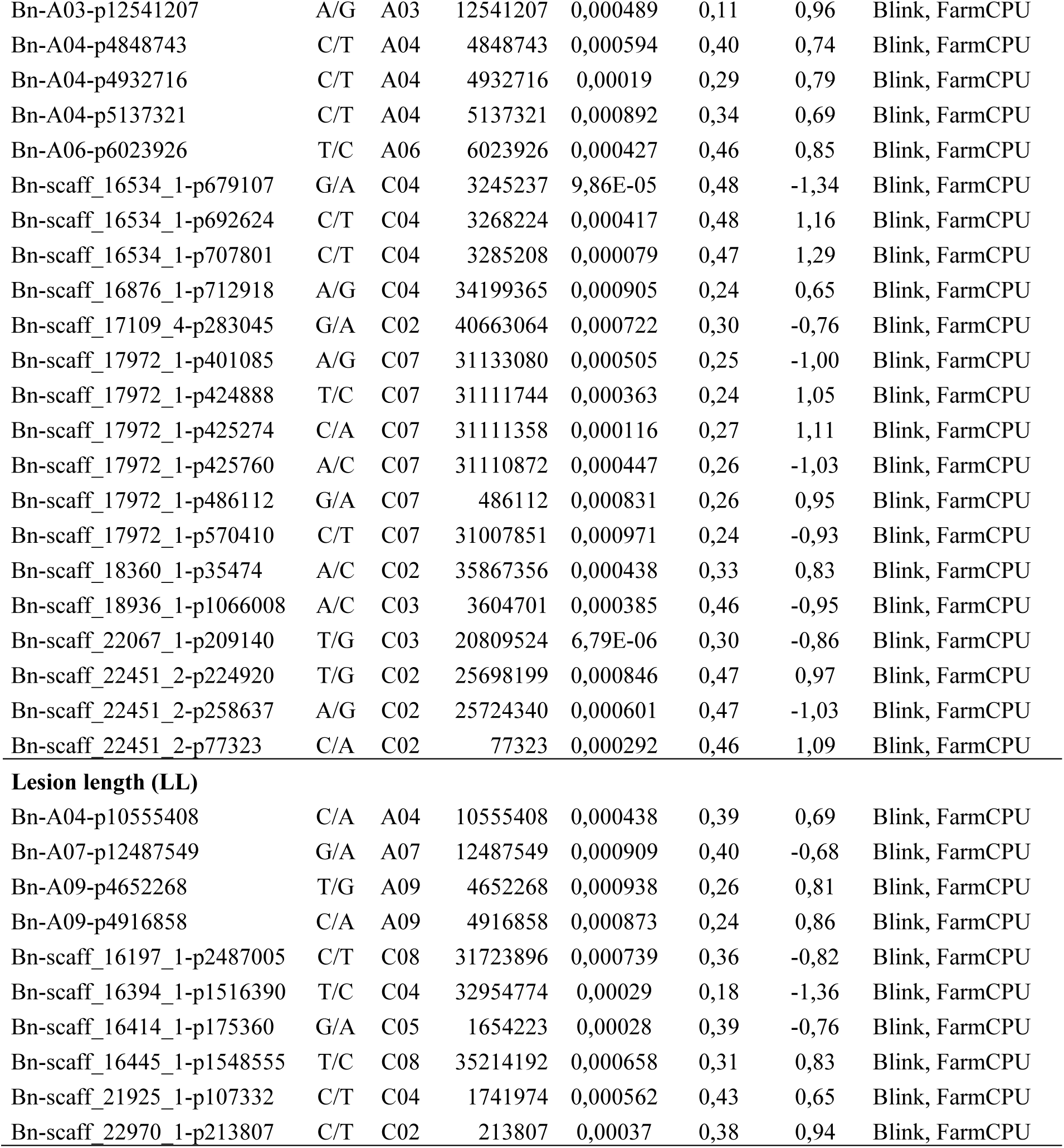
MTAs linked to Sclerotinia stem rot (SSR) resistance in oilseed rape accessions.

| SNP | Allele | Chr | Pos | *P. value* | MAF | Effect | Model |
| --- | --- | --- | --- | --- | --- | --- | --- |
| **Relative lesion area (RLA)** |  |  |  |  |  |  |  |
| Bn-A01-p19174115 | G/A | A01 | 19174115 | 0,00058 | 0,46 | -0,02 | Blink, FarmCPU |
| Bn-A01-p5265598 | T/C | A01 | 5265598 | 0,000525 | 0,44 | 0,02 | Blink, FarmCPU |
| Bn-A01-p5269368 | A/G | A01 | 5269368 | 0,000341 | 0,46 | -0,02 | Blink, FarmCPU |
| Bn-A03-p10027020 | C/A | A03 | 10027020 | 0,000838 | 0,30 | -0,02 | Blink, FarmCPU |
| Bn-A06-p1015217 | A/G | A06 | 1015217 | 0,000837 | 0,5 | -0,02 | Blink, FarmCPU |
| Bn-A07-p14766907 | G/T | A07 | 14766907 | 0,000241 | 0,35 | -0,02 | Blink, FarmCPU |
| Bn-A07-p14768265 | C/T | A07 | 14768265 | 0,000241 | 0,35 | -0,02 | Blink, FarmCPU |
| Bn-A07-p14770619 | C/A | A07 | 14770619 | 0,00043 | 0,34 | 0,02 | Blink, FarmCPU |
| Bn-A07-p14792267 | C/A | A07 | 14792267 | 0,000161 | 0,36 | 0,02 | Blink, FarmCPU |
| Bn-A07-p15165145 | C/T | A07 | 15165145 | 0,000262 | 0,35 | -0,02 | Blink, FarmCPU |
| Bn-A07-p17388825 | C/T | A07 | 17388825 | 0,000918 | 0,09 | -0,03 | Blink, FarmCPU |
| **Lesion Area (LA)** |  |  |  |  |  |  |  |
| Bn-A01-p2291940 | C/T | A01 | 2291940 | 0,000155 | 0,48 | -0,81 | Blink, FarmCPU |
| Bn-A01-p24575577 | T/C | A01 | 24575577 | 6,58E-05 | 0,12 | -1,00 | Blink, FarmCPU |
| Bn-A01-p4353657 | T/C | A01 | 4353657 | 0,000235 | 0,35 | 0,97 | Blink, FarmCPU |
| Bn-A02-p23835683 | C/A | A02 | 23835683 | 0,000815 | 0,42 | -0,82 | Blink, FarmCPU |
| Bn-A03-p12541207 | A/G | A03 | 12541207 | 0,000489 | 0,11 | 0,96 | Blink, FarmCPU |
| Bn-A04-p4848743 | C/T | A04 | 4848743 | 0,000594 | 0,40 | 0,74 | Blink, FarmCPU |
| Bn-A04-p4932716 | C/T | A04 | 4932716 | 0,00019 | 0,29 | 0,79 | Blink, FarmCPU |
| Bn-A04-p5137321 | C/T | A04 | 5137321 | 0,000892 | 0,34 | 0,69 | Blink, FarmCPU |
| Bn-A06-p6023926 | T/C | A06 | 6023926 | 0,000427 | 0,46 | 0,85 | Blink, FarmCPU |
| Bn-scaff_16534_1-p679107 | G/A | C04 | 3245237 | 9,86E-05 | 0,48 | -1,34 | Blink, FarmCPU |
| Bn-scaff_16534_1-p692624 | C/T | C04 | 3268224 | 0,000417 | 0,48 | 1,16 | Blink, FarmCPU |
| Bn-scaff_16534_1-p707801 | C/T | C04 | 3285208 | 0,000079 | 0,47 | 1,29 | Blink, FarmCPU |
| Bn-scaff_16876_1-p712918 | A/G | C04 | 34199365 | 0,000905 | 0,24 | 0,65 | Blink, FarmCPU |
| Bn-scaff_17109_4-p283045 | G/A | C02 | 40663064 | 0,000722 | 0,30 | -0,76 | Blink, FarmCPU |
| Bn-scaff_17972_1-p401085 | A/G | C07 | 31133080 | 0,000505 | 0,25 | -1,00 | Blink, FarmCPU |
| Bn-scaff_17972_1-p424888 | T/C | C07 | 31111744 | 0,000363 | 0,24 | 1,05 | Blink, FarmCPU |
| Bn-scaff_17972_1-p425274 | C/A | C07 | 31111358 | 0,000116 | 0,27 | 1,11 | Blink, FarmCPU |
| Bn-scaff_17972_1-p425760 | A/C | C07 | 31110872 | 0,000447 | 0,26 | -1,03 | Blink, FarmCPU |
| Bn-scaff_17972_1-p486112 | G/A | C07 | 486112 | 0,000831 | 0,26 | 0,95 | Blink, FarmCPU |
| Bn-scaff_17972_1-p570410 | C/T | C07 | 31007851 | 0,000971 | 0,24 | -0,93 | Blink, FarmCPU |
| Bn-scaff_18360_1-p35474 | A/C | C02 | 35867356 | 0,000438 | 0,33 | 0,83 | Blink, FarmCPU |
| Bn-scaff_18936_1-p1066008 | A/C | C03 | 3604701 | 0,000385 | 0,46 | -0,95 | Blink, FarmCPU |
| Bn-scaff_22067_1-p209140 | T/G | C03 | 20809524 | 6,79E-06 | 0,30 | -0,86 | Blink, FarmCPU |
| Bn-scaff_22451_2-p224920 | T/G | C02 | 25698199 | 0,000846 | 0,47 | 0,97 | Blink, FarmCPU |
| Bn-scaff_22451_2-p258637 | A/G | C02 | 25724340 | 0,000601 | 0,47 | -1,03 | Blink, FarmCPU |
| Bn-scaff_22451_2-p77323 | C/A | C02 | 77323 | 0,000292 | 0,46 | 1,09 | Blink, FarmCPU |
| Lesion length (LL) |  |  |  |  |  |  |  |
| Bn-A04-p10555408 | C/A | A04 | 10555408 | 0,000438 | 0,39 | 0,69 | Blink, FarmCPU |
| Bn-A07-p12487549 | G/A | A07 | 12487549 | 0,000909 | 0,40 | -0,68 | Blink, FarmCPU |
| Bn-A09-p4652268 | T/G | A09 | 4652268 | 0,000938 | 0,26 | 0,81 | Blink, FarmCPU |
| Bn-A09-p4916858 | C/A | A09 | 4916858 | 0,000873 | 0,24 | 0,86 | Blink, FarmCPU |
| Bn-scaff_16197_1-p2487005 | C/T | C08 | 31723896 | 0,000739 | 0,36 | -0,82 | Blink, FarmCPU |
| Bn-scaff_16394_1-p1516390 | T/C | C04 | 32954774 | 0,00029 | 0,18 | -1,36 | Blink, FarmCPU |
| Bn-scaff_16414_1-p175360 | G/A | C05 | 1654223 | 0,00028 | 0,39 | -0,76 | Blink, FarmCPU |
| Bn-scaff_16445_1-p1548555 | T/C | C08 | 35214192 | 0,000658 | 0,31 | 0,83 | Blink, FarmCPU |
| Bn-scaff_21925_1-p107332 | C/T | C04 | 1741974 | 0,000562 | 0,43 | 0,65 | Blink, FarmCPU |
| Bn-scaff_22970_1-p213807 | C/T | C02 | 213807 | 0,00037 | 0,38 | 0,94 | Blink, FarmCPU |

### MTAs for Chlorophyll content in SSR-infected oilseed rape

The GWAS analysis identified 24 MTAs significantly linked to the chlorophyll traits; seven for SPAD before inoculation (SPADH), six for SPAD at three days post inoculation (SPADI), and 11 MTAs for chlorophyll index (CI) (Table 4 and Figure 4). All the MTAs for SPADH and six for SPADI were identified in both FarmCPU and Blink models, while for CSI six MTAs were identified in the FarmCPU and four were identified in the Blink model. SNP markers Bn-scaff_20461_1-p249973 and Bn-scaff_20461_1-p178421 located on chromosome C02 and Bn-scaff_16361_1-p1343410 located on chromosome C08 were closely linked to CSI and SPADI (Table 4).

**Figure 4.**
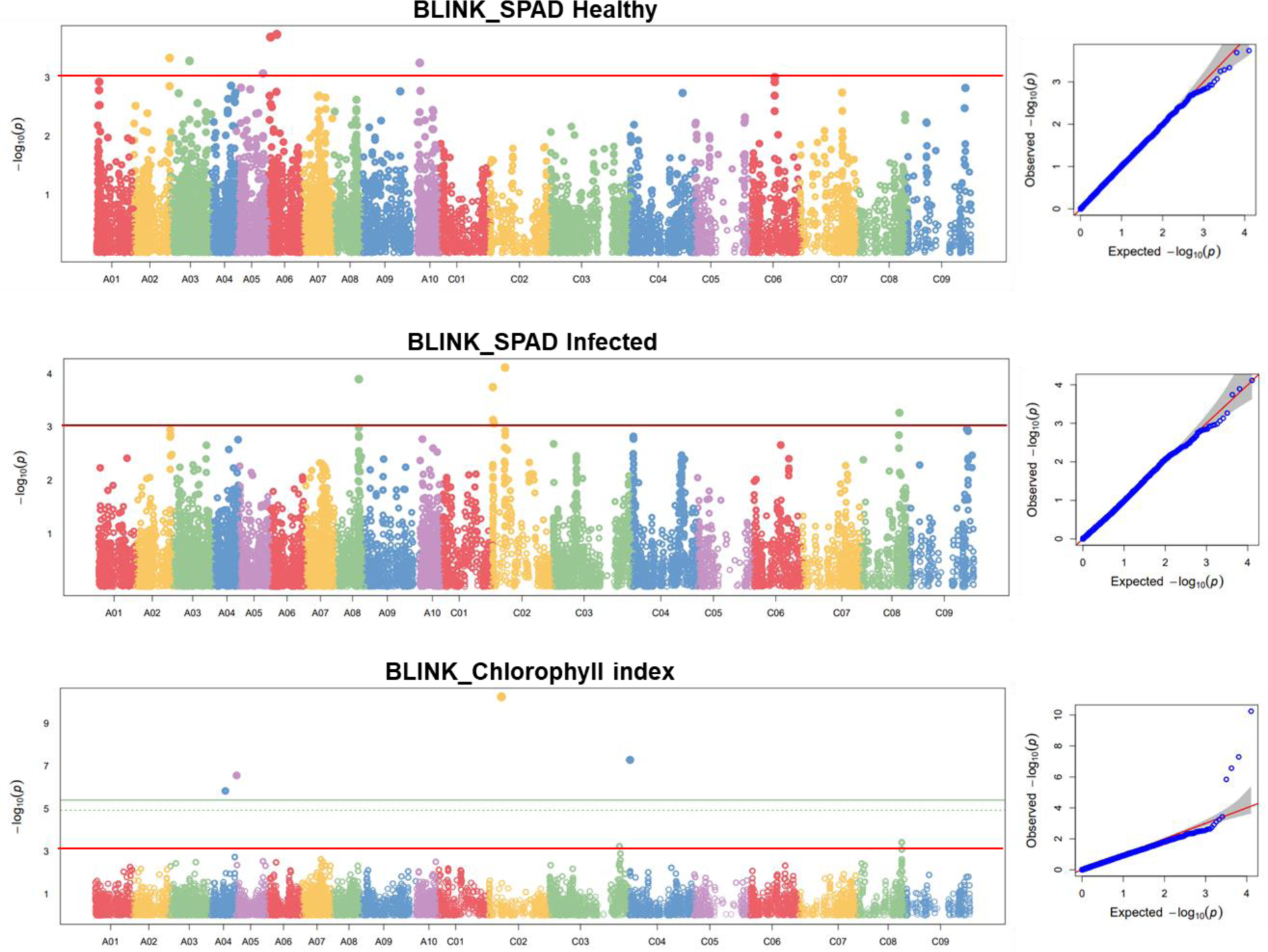
Manhattan (left) and Q-Q (right) plots for the GWAS for the SPAD measurements in healthy leaves before inoculation (SPADH), at 3 days post inoculation (SPADI) with Sclerotinia stem rot (SSR), and the chlorophyll index (CI) in oilseed rape accessions.

Six of the seven MTAs linked to the SPADH were identified in subgenome A including two MTAs located on chromosome A06 while there was only one MTA that was located on C06. Conversely, four of the six MTAs significantly associated with SPADI were identified in the C subgenome, Bn-scaff_20461_1-p249973 (9.42 Mb) and Bn-scaff_20942_1-p1093417 (10.93 Mb) as well as Bn-scaff_20461_1-p178421 (0.18 Mb) and Bn-scaff_15712_5-p313157 (0.31 Mb) on chromosome C02 (Table 4).

For CI, four MTAs were on chromosome C02, three on C02, and one on each subgenome A chromosomes A02, A04, and A05. Three of the four MTAs identified on C02 were located on 9.42Mb (Bn-scaff_20461_1-p249973), 9.55Mb (Bn-scaff_20461_1-p110039), and 9.56Mb (Bn-scaff_20461_1-p99520) (Table 5). The three are within the LD region for chromosome C02 indicating the presence of QTL region on this chromosome. Likewise, the three MTAs for CI identified on C08 at 28.81Mb (Bn-scaff_16361_1-p1343410), 28.41Mb (Bn-scaff_16361_1-p935272), and 33.36Mb (Bn-scaff_16197_1-p659483) (Table 4). These SNPs within the LD score indicate they are linked.

**Table 4.** MTAs linked to chlorophyll content and stability in oilseed rape accessions.

| SNP | Allele | Chr | Pos | *P. value* | MAF | Effect | Model |
| --- | --- | --- | --- | --- | --- | --- | --- |
| **Chlorophyll Index (CI)** |  |  |  |  |  |  |  |
| Bn-A02-p26154951 | A/G | A02 | 26154951 | 0,000494 | 0,17 | -1,79 | FarmCPU |
| Bn-A04-p11329756 | T/G | A04 | 11329756 | 1,46E-06 | 0,26 | 1,82 | Blink |
| Bn-A05-p967131 | C/T | A05 | 967131 | 2,74E-07 | 0,10 | -2,11 | Blink |
| Bn-scaff_20461_1-p249973 | A/G | C02 | 9420451 | 5,73E-11 | 0,48 | -2,39 | Blink, FarmCPU |
| Bn-scaff_20461_1-p178421 | T/C | C02 | 178421 | 0,000167 | 0,43 | 1,94 | FarmCPU |
| Bn-scaff_20461_1-p99520 | A/G | C02 | 9562691 | 0,000267 | 0,49 | -2,06 | FarmCPU |
| Bn-scaff_20461_1-p110039 | G/A | C02 | 9550142 | 0,000562 | 0,46 | 1,83 | FarmCPU |
| Bn-scaff_15832_1-p52737 | C/A | C04 | 378587 | 5,12E-08 | 0,37 | -1,41 | Blink |
| Bn-scaff_16361_1-p1343410 | A/G | C08 | 28813035 | 0,000303 | 0,48 | 1,52 | FarmCPU |
| Bn-scaff_16361_1-p935272 | G/A | C08 | 28413763 | 0,000357 | 0,43 | 1,50 | FarmCPU |
| Bn-scaff_16197_1-p659483 | T/C | C08 | 33364657 | 0,000376 | 0,26 | -0,99 | Blink |
| **SPAD Healthy (SPADH)** |  |  |  |  |  |  |  |
| Bn-A02-p27203061 | G/A | A02 | 27203061 | 0,000463 | 0,49 | 0,51 | Blink, FarmCPU |
| Bn-A03-p14583041 | T/G | A03 | 14583041 | 0,000521 | 0,48 | -0,61 | Blink, FarmCPU |
| Bn-A05-p20141883 | T/G | A05 | 20141883 | 0,000855 | 0,40 | -0,59 | Blink, FarmCPU |
| Bn-A06-p6439396 | G/A | A06 | 6439396 | 0,000184 | 0,26 | 0,87 | Blink, FarmCPU |
| Bn-A06-p1768254 | G/A | A06 | 1768254 | 0,000204 | 0,40 | 0,60 | Blink, FarmCPU |
| Bn-A10-p2268881 | G/A | A10 | 2268881 | 0,000562 | 0,26 | 0,79 | Blink, FarmCPU |
| Bn-scaff_15818_2-p1213540 | A/G | C06 | 17603819 | 0,000992 | 0,18 | -0,67 | Blink, FarmCPU |
| **SPAD Infected (SPADI)** |  |  |  |  |  |  |  |
| Bn-A08-p16859028 | A/G | A08 | 16859028 | 0,000128 | 0,36 | 0,89 | Blink, FarmCPU |
| Bn-scaff_20461_1-p249973 | A/G | C02 | 9420451 | 7,71E-05 | 0,48 | -1,16 | Blink, FarmCPU |
| Bn-scaff_20461_1-p178421 | T/C | C02 | 178421 | 0,00018 | 0,43 | 1,08 | Blink, FarmCPU |
| Bn-scaff_15712_5-p313157 | A/G | C02 | 313157 | 0,000738 | 0,42 | -1,178 | Blink, FarmCPU |
| Bn-scaff_20942_1-p1093417 | C/T | C02 | 1093417 | 0,000865 | 0,45 | -1,15 | Blink, FarmCPU |
| Bn-scaff_16361_1-p1343410 | A/G | C08 | 28813035 | 0,000543 | 0,48 | 0,82 | Blink, FarmCPU |

### Phyllosphere Microbiota diversity of rapeseed in resistance and susceptible accessions

This study used 16S rRNA gene sequencing to elucidate the diversity and composition of microbial (bacterial and fungal) communities within the phyllosphere of resistant and susceptible rapeseed varieties and understand their potential role in plant resistance mechanisms.

#### Bacterial Communities

Our analysis revealed significant differences in the bacterial alpha diversity between the resistant and susceptible rapeseed varieties (Figure 5A). The bacterial richness, as indicated by the Chao1 estimator, was significantly higher in susceptible varieties (Wilcoxson test, p-value < 0.005). Additionally, the Shannon index, which accounts for both richness and evenness, also showed significantly higher overall diversity in the susceptible varieties (Wilcoxson test, p-value < 0.005). These results suggest that the phyllosphere of susceptible rapeseed varieties harbors more diverse communities. In contrast, the fungal microbiota diversity did not exhibit significant differences between the resistant and susceptible varieties (Figure 5B, Wilcoxson test, p-value > 0.005).

**Figure 5.**
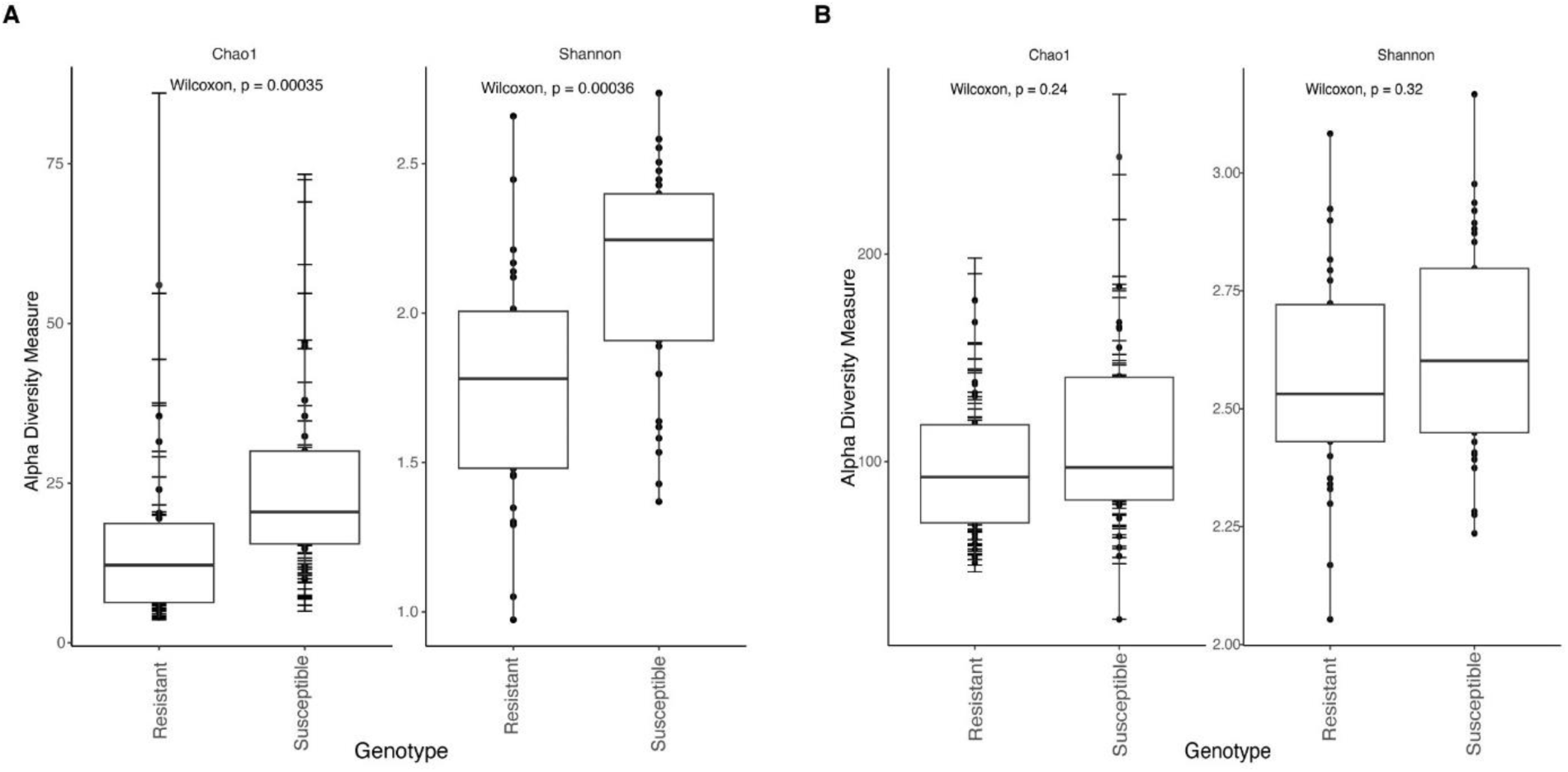
Microbial community analysis: Bacterial (A) and fungal (B) communities alpha diversity measures in resistant and susceptible rapeseed varieties using Chao1 and Shannon indexes. Statistical significance was determined using the Wilcoxson test.

### Taxonomic structure and relative abundance of the dominant microbial communities

We identified 18 bacterial and four fungal phyla within the rapeseed microbiome. Proteobacteria was the most prevalent bacterial phylum in both resistant and susceptible samples, with a relative abundance ranging from 52% in susceptible and 58% in resistant varieties (Figure 6A and Supplementary Table S2). The second most prevalent phylum was *Actinobacteria,* constituting 21% of the relative abundance in both varieties. *Firmicutes* represented approximately 13% of the bacterial community (Supplementary Table S2). Additionally, *Bacteroidota* contributed to approximately 5-6% of the total relative abundance (Figure 6A). In the fungal community, the phylum *Ascomycota* was the most abundant in resistant and susceptible samples, with relative abundances ranging from 58% to 64% (Figure 6B and Supplementary Table S3). This was followed by *fungi-phy-incertae sedis*, constituting 31% to 33% of the fungal community (Supplementary Table S3). The phylum *Basidiomycota* contributed 4% to 7% in resistant and susceptible samples (Figure 6B).

**Figure 6.**
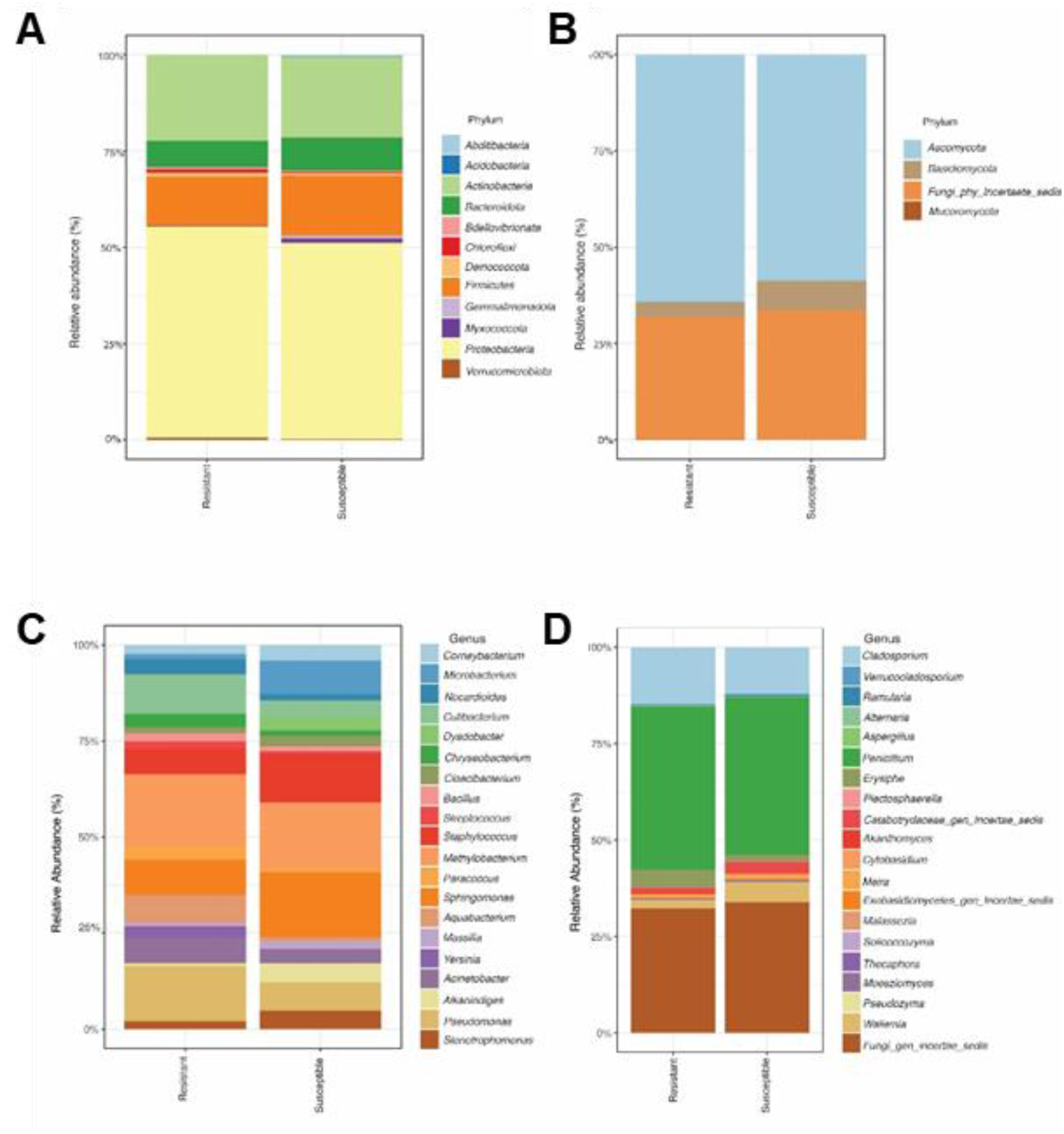
Community composition of the resistant and susceptible rapeseed varieties using the relative abundance at Phylum and genus levels. The relative abundance of the most abundant phyla in Bacteria (A) and Fungi (B), followed by the relative abundance of the top 20 bacterial (C) and fungal (D) genera, were visualized in the bar chart.

The most abundant bacterial genera were *Pseudomonas*, *Methylobacterium*, *Sphingomonas*, *Staphylococcus*, *Microbacterium,* and *Aquabacterium* (Figure 6C). Genera such as *Pseudomonas*, *Methylobacterium*, *Aquabacterium*, *Yersinia*, and *Cutibacterium* were more prevalent in resistant samples (Figure 6C and Supplementary Table S4). In contrast, *Sphingomonas*, *Staphylococcus*, and *Microbacterium* were more abundant in susceptible samples (Figure 6C). Moreover, the linear discriminant analysis (LDA) revealed the specific taxa responsible for the observed significant differences in the composition of resistant and susceptible microbiomes (Figure 7A). 5 taxa were significantly enriched in resistant varieties and had an LDA score of > 4. The most enriched taxa were *Pseudomonas (genus), Pseudomonadales (*Order*), Pseudomonadaceae* (Family), and *Pseudomonas* (Species). 9 taxa significantly differed in susceptible rapeseed varieties. The members of the *Micrococcales* (order) had a larger size effect with an LDA score > 4, followed by *Sphingobactriales* and *Massilia* (Figure 7A).

**Figure 7.**
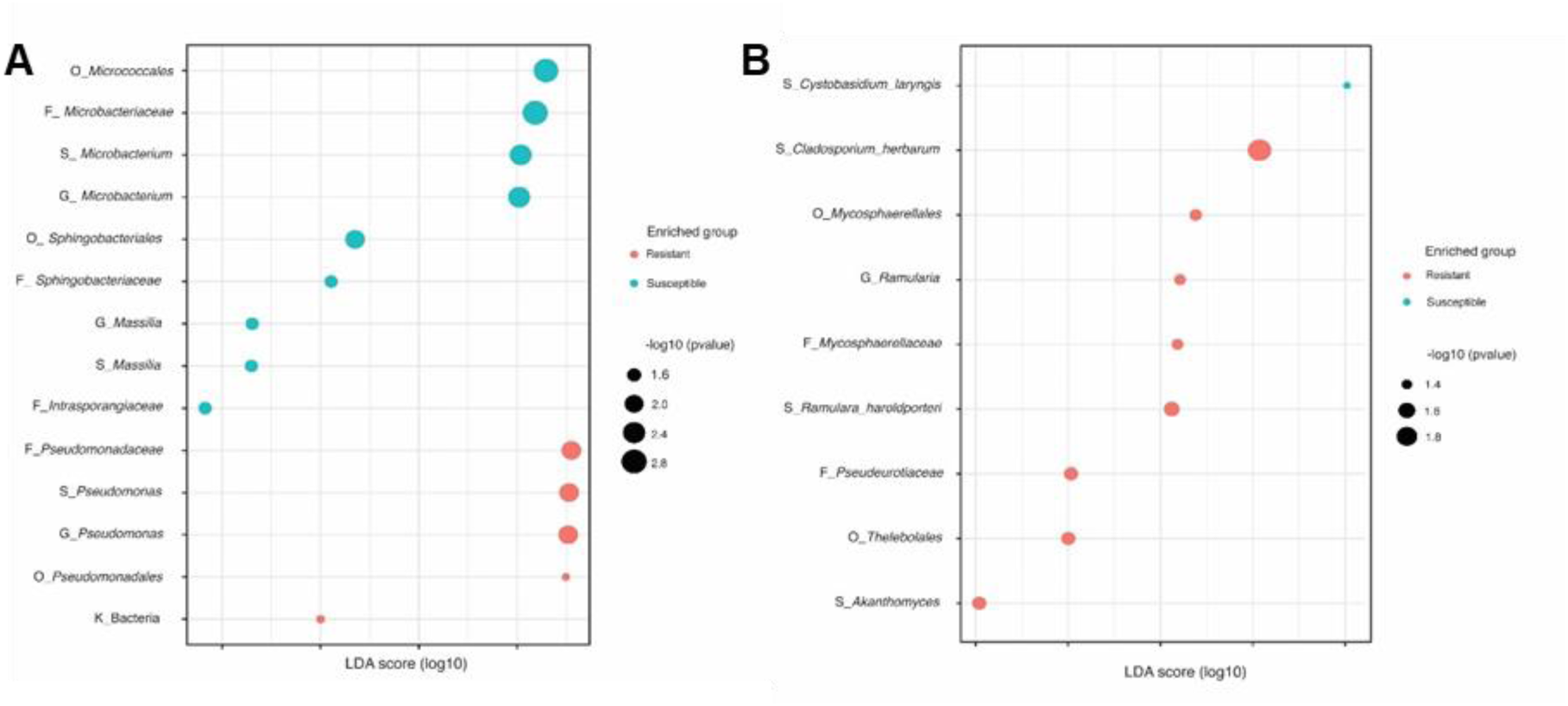
Differential abundance analysis using linear discriminant analysis (LDA). This figure shows the differentially abundant bacterial taxa (A), and fungal taxa (B) identified in resistant and susceptible oilseed rape accessions.

**Figure 8.**
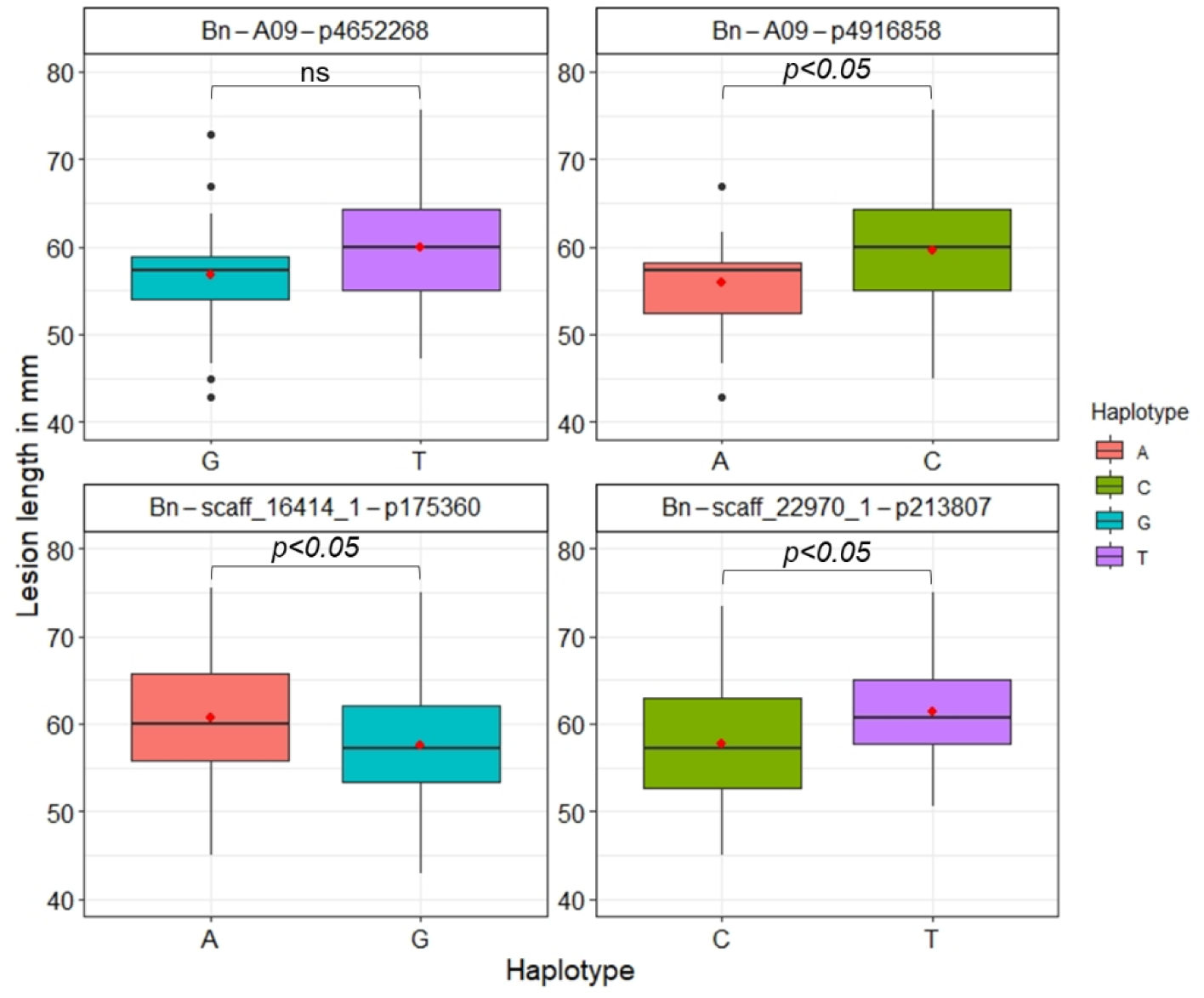
Contribution of the favorable and alternative alleles of the MTAs for the lesion length variation among the oilseed rape accessions.

The three most abundant genera across all samples for the fungal community were *Penicillium*, *Cladosporium*, and genera classified as *fungi_gen_ Incertae_sedis* (Figure 6D and Supplementary Table S5). Although the relative abundance of these fungal genera remained relatively stable between resistant and susceptible samples, LDA showed that resistant varieties had nine enriched taxa. In comparison, susceptible varieties had only one (*Cystobasidium* _*laryngis)* differentially enriched taxa (Figure 7B). The most enriched taxa in resistant varieties were members of the *Mycosphaerellales*, *Thelebolales*, and *Akanthomyces* (Figure 7B).

### SSR-related variants contributing to SSR resistance and microbial recruitment

The most relatively susceptible and resistant accessions of the oilseed rape germplasm were used for investigating the microbiome recruitment variation between these groups. The resistant group had relatively lower lesion length (LL), lesion area (LA), and relative lesion area (RLA) compared to the susceptible group (Supplementary Figure 1A). However, there was no significant difference between the resistant and susceptible groups for the SPADH, SPADI, and CI values (Supplementary Figures 1B). The allelic variations, in the 47 MTAs detected linked to SSR resistance (Table 5), between these resistance and susceptible groups are presented (Supplementary Table S6). The most consistent allele variation between resistance and susceptible groups was MTAs associated with the lesion length (Table 6). The top four MTAs with the most significant phenotypic effect were Bn-A09-p4652268 (T/G) and Bn-A09-p4916858 (C/A) located on A09, and Bn-scaff_22970_1-p213807 (C/T) and Bn-scaff_16414_1-p175360 (G/A) located on C02 and C05 respectively (Table 6). In the whole oilseed rape germplasm investigated, the mean lesion length value for the favorable allele, found commonly in resistant accessions, was smaller compared to the alternative allele commonly found in the susceptible group (Figure 6). The favorable alleles found on the A09 were rare in the germplasm while those found in the C02 and C05 were common, compared to their respective alternative alleles (Figure 9).

**Table 6.**
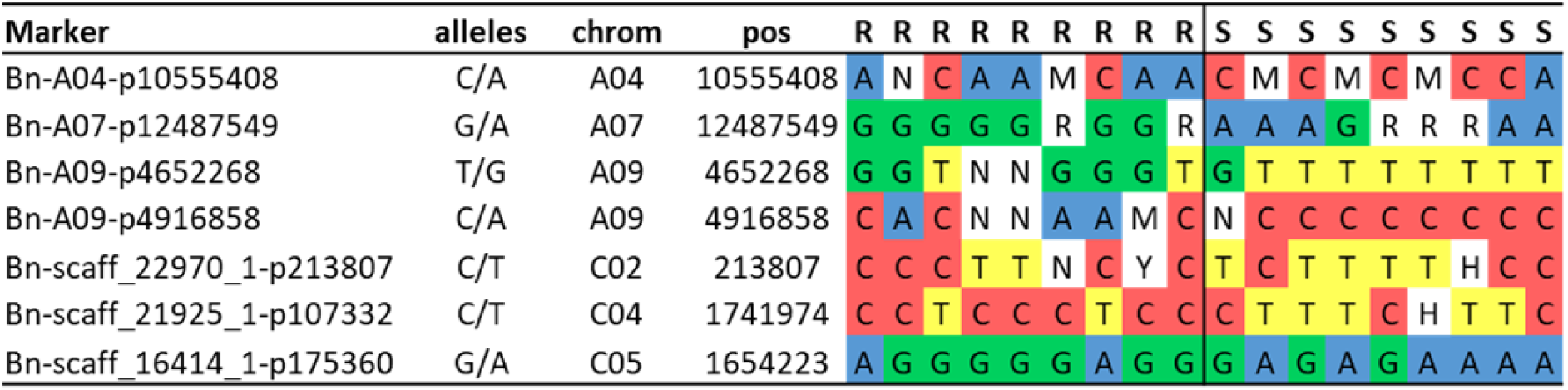
Allele variation between the most resistant and susceptible accessions.

## Discussion

Identifying resistance genotypes within a diverse germplasm and detecting the genetic basis for resistance is critical for enhancing host resistance and developing resistant varieties of economically important crops. This is particularly important to mitigate the impact of Sclerotinia stem rot on oilseed rape production as cultural control practices are ineffective and laborious and it is difficult to predict the optimal time for applying fungicides (Derbyshire and Denton-Giles, 2016). However, no genotype of rape seed with complete resistance against the pathogen has been found so far (Neik et al., 2017; Ding et al., 2021). Given population structure and aggressiveness and rich effector repertoire diversity of the pathogen diversity (Taylor et al., 2015; Yu et al., 2020; Gupta et al., 2022) as well as resistance variation among subpopulations of oilseed rape (Neik et al., 2017), investigating the resistance in locally adapted germplasm would provide resistance sources for breeding programs targeted at particular regions. Enhancing the level of resistance in oilseed rape through the pyramiding existing partial resistance is challenging because the QTLs for such resistance explain only a low percentage of the phenotypic variations and lack reproducibility (Ding et al., 2021). Hence, beyond the genetics approach, understanding agronomic and physiological processes contributing to enhanced levels of the existing partial resistance in oilseed rape is needed to limit the impact of the disease.

### Sclerotinia stem rot resistance in rape seed accessions

In total, 47 MTAs spanning across all chromosomes except in A10, C01, and C06 were detected associated with the SSR resistance traits LL, LA, and RLA. Twenty-four of these MTAs were from the A-subgenome and 26 from the C-subgenome of the crop, with chromosomes A07 and C07 contributing seven and six MTAs. This is in agreement with previous findings which showed that SSR resistance is a complex trait controlled by several QTLs with minor effects (Wei et al., 2014; Wu et al., 2016; Roy et al., 2021). The findings indicate both subgenomes play a crucial role in the SSR resistance of oilseed rape. A different set of significant SNPs were identified for Stem lesion length and Stem lesion width (Roy et al., 2021; Roy et al., 2024) and 17 SNPs for stem lesion length (Wei et al., 2016) have been previously identified in oilseed rape. These studies were based on field experiments and stem tissue, but the results suggest the possibility of finding different SNPs for SSR resistance traits LL, LA, and RLA found in the current study. Analyzing several aspects of the Sclerotinia stem rot disease growth on plant tissue and crucial for establishing a relationship between the disease symptoms and genetic bases regulating genotype-specific resistance response to the pathogen. However, the MTAs detected in this study had minimal effect on the phenotypes measured compared to the previous studies (Wei et al., 2016; Roy et al., 2021; Roy et al., 2024). This could be due to less number of accessions investigated in the present study, and the previous studies were focused on stem resistance, conducted in field conditions hence microclimate factors can influence SSR disease development and plant response (Ficke et al., 2018). However, all the significant SNPs associated with SSR resistance were detected using the most stringent models Blink (Huang et al., 2018) and FarmCPU models (Liu et al., 2016), which shows the relevance of the MTAs under the current study conditions. Indeed, we found SNPs consistently differ between the relatively resistant and susceptible groups, with the favorable alleles showing lower LL, LA, and RLA compared to the alternative alleles. Particularly the SNPs linked to LL Bn-A09-p4652268, Bn-A09-p4916858, Bn-scaff_22970_1-p213807, Bn-scaff_16414_1-p175360 were interesting as the alleles segregate in the most relatively resistant and susceptible accessions used for the microbial recruitment investigation.

### Chlorophyll level during the Sclerotinia stem rot infection

SSR infection reduces the leaf area for photosynthesis hence maintaining high chlorophyll content is crucial for producing photosynthate and reducing yield loss due to the pathogen infection. A total of 7 MTAs were detected for SPADH, 6 MTAs for SPADI, and 11 MTAs for CI. Of these MTAs, Bn-scaff_20461_1-p249973 and Bn-scaff_20461_1-p178421 located on C02, and Bn-scaff_16361_1-p1343410 located on C08 were associated with SPADI and CI. CI is an important trait because the SSR is a necrotrophic fungus (Gupta et al., 2022) reduces the leaf area hence the SPADI. Considering this, CI is an important trait in maintaining the chlorophyll levels and the photosynthetic rate. Chlorophyll indices could be used for deteting plant diseases (Odilbekov et al., 2018; Koc et al., 2022). In agreement with this study, a significant majority of the 281 single-nucleotide polymorphisms (SNPs) associated with photosynthetic pigment (Xu et al., 2022) and ten out of fourteen SNPs linked to chlorophyll synthesis (Hu et al., 2022) in oilseed rape were located in the C-subgenome. Moreover, a chlorophyll-deficient dominant locus (CDE1) in oilseed rape was mapped to C08 (Wang et al., 2016). The findings indicate the C-subgenome’s crucial role in maintaining the chlorophyll content and photosynthesis rate, and both traits should be considered as an SSR resistance-related trait in oilseed rape.

### Microbial recruitment in sclerotinia-resistant and susceptible oilseed rape accessions

The composition and structure of the phyllosphere are critical to its impact on the host plant, highlighting the need to understand the mechanisms behind its assembly and dynamics (Hawkes et al., 2021). Plant genotype is a crucial factor influencing the microbial diversity of the phyllosphere. Our study investigates the microbiota of resistant and susceptible rapeseed varieties, focusing on bacterial and fungal communities. We found significant differences in bacterial diversity between the two genotypes, while fungal communities remain unaffected by these variations.

The genetic variations between resistant and susceptible rapeseed varieties notably influence bacterial diversity. The observed higher alpha diversity in the susceptible rapeseed varieties than resistant ones could be due to several factors. For instance, susceptible varieties are more prone to pathogen infection, leading to severe tissue damage and a weaker immune system. This environment allows a broader range of microbes to coexist, including opportunistic pathogens and commensals. Moreover, the compromised defense mechanism in susceptible plants leads to frequent invasions by environmental microbes, contributing to higher microbial diversity. Conversely, the robust immune system in the resistant varieties suppresses the invasion of certain microbes, selectively enriching the specific microbial communities and resulting in less diverse but specialized microbial communities.

In this study, this selective enrichment is more evident by the significantly enriched members of the *Pseudomonas* in resistant rapeseed varieties. Genus *Pseudomonas* is widely known for its role in plant growth promotion and biological control of plant diseases. These bacteria are ubiquitous in nature and are characterized by their versatile metabolic activities. Strains and species of this genus enhance plant growth by facilitating nutrient acquisition, regulating plant hormone levels, and inducing a systemic response in plants through disease suppression (biological control). They achieve this via the production of antibiotics and the regulation of signaling molecules (Kumar et al., 2017; Jegan et al., 2018). Additionally, resistant varieties enriched the phyllosphere-abundant genus *Methylobacterium*, which is known to promote plant growth and development (Dourado et al., 2015).

Moreover, fungal communities significantly enriched in resistant varieties such as Thelebolales and Pseuderotiaceae are known to produce secondary metabolites, antifreeze proteins and ice-freezing proteins, which strengthen the resistance rapeseed microbiome. Besides, the species *Akanthomyces* ensures protection against a wide array of pests and, including antifungal and bacterial activity against *Sclerotinia sclerotiorum*, *Rhizoctonia solani*, *Aspergillus flavus* and *Staphylococcus aureus* pathogens (Gurulingappa et al., 2010; Dash et al., 2018; Nicoletti and Becchimanzi, 2020).

On the other hand, susceptible rapeseed varieties exhibited higher abundances of genera associated with stress tolerance (*Sphingomonas*) (Asaf et al., 2020) and plant growth and disease control (*Stenotrophomonas* and *Microbacterium*) (Kumar et al., 2023). Bacterial species from the genera *Microbacterium* and *Stenotrophomonas* were identified at blast lesion, which reduced the blast severity and promoted plant growth (Behrendt et al., 2001; Md Gulzar and Mazumder, 2023). These findings suggest an increased abundance of specific microbial taxa in different genotypes could recruit specific ones to fulfill their demands. This implies that genetic differences in rapeseed varieties may drive selective enrichment of certain bacterial taxa’s relative abundance and recruiting specific microbes rather than exclude them. Mainly, phyla Proteobacteria and Bacteroidota were the most responsive to these genotype variations, similar to previous reports (Taye et al., 2020).

Unlike bacterial communities, fungal communities exhibited stable diversity despite minor changes in the relative abundance of dominant taxa. This stability indicates that fungal assemblages are less influenced by the host plant’s genetic variations and more affected by factors such as geographical location and environmental stress (Barret et al., 2015; Barret et al., 2016). This stable fungal community structure across resistant and susceptible varieties suggests that the phyllosphere fungal composition does not directly influence the resistance function observed in these rapeseed varieties.

Overall, this study highlights the impact of genetic variation on the dynamics of the phyllosphere microbial community. The higher relative dominance of specific microbial taxa supports the notion that plants actively select particular microbial communities to meet their requirements. These findings imply that breeding for specific resistance traits in oilseed rape could indirectly influence the composition and function of the phyllosphere microbiome. These findings contribute to a better understanding of the relationships between plant genotypes and their associated microbiomes, with potential implications for agricultural practices and crop improvement strategies.

## Conclusion

The study provided the genetic regions playing a crucial role in Sclerotinia stem rot resistance and maintaining the chlorophyll content during the infection. As the pathogen expands on the leaf and the photosynthetic efficient area reduces hence increasing or maintaining higher chlorophyll content on the uninfected part of the leaf could be a tolerance mechanism in oilseed rape for reducing the impact of the pathogen on yield and quality. Importantly, the SSR resistance level and chlorophyll index differences between the resistant and susceptible accessions were reflected in the microbe composition. The study has limitations related to the size of the GWAS panel and only one strain of Sclerotinia was used for resistance screening. However, the study provides insights into multifaceted strategies and their genetic basis for applying breeding techniques to improve the resistance and further enhance the resistance level by taking into consideration the genetic basis of the resistance phenotype, chlorophyll index, and microbial recruitment in oilseed rape.

## Acknowledgments

KBA acknowledges support from KSLA (GFS2021-0082).

## Contributions

AC, KBA, FO, and RV conceived and initiated the project. KBA, VT, SS, and AC conducted resistance screening and genotyping of accessions. FG and RV conducted the microbiome composition experiment. KBA and SG analyzed the GWAS data and microbiome data, respectively. KBA prepared the manuscript, assisted by SM, AC, and SG. All authors read the final version of the manuscript and approved its publication.

## Supplementary Table Legends

Supplementary Table S1. List of oilseed rape accessions used for GWAS analysis.

Supplementary Table S2. Relative abundance (%) of each bacterial amplicon sequence variant (ASV) across the resistant and susceptible rapeseed varieties at the phylum level.

Supplementary Table S3. Relative abundance (%) of each fungal amplicon sequence variant (ASV) across the resistant and susceptible rapeseed varieties at the phylum level.

Supplementary Table S4. Relative abundance (%) of each bacterial amplicon sequence variant (ASV) across the resistant and susceptible rapeseed varieties at the genus level.

Supplementary Table S5. Relative abundance (%) of each fungal amplicon sequence variant (ASV) across the resistant and susceptible rapeseed varieties at the genus level.

Supplementary Table S6. Haplotype variation, on the SNPs linked to SSR resistance, between the most resistant (R) and susceptible (S) group of oilseed rape accessions.

