## Supplementary Figure 1 for "Genomic loci for sclerotinia stem rot resistance and chlorophyll stability in *Brassica napus*: integrating GWAS with microbiome insights"

### Supplementary Figures

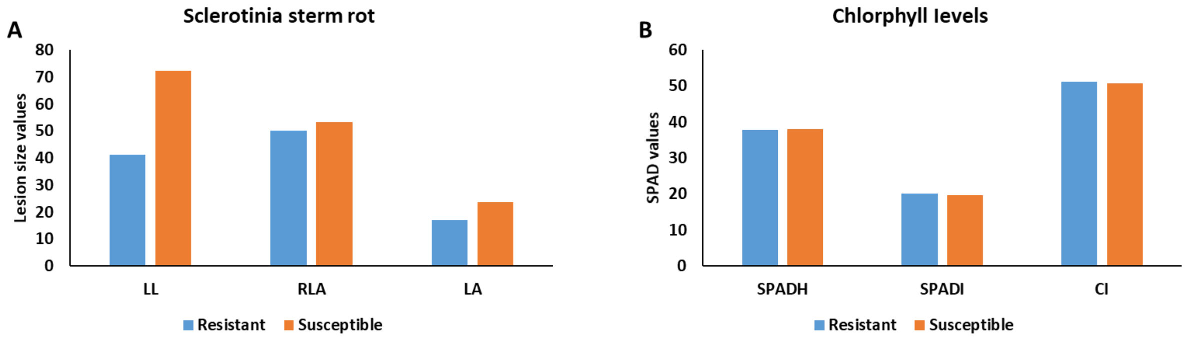


**Supplementary Figure 1**. Sclerotinia stem rot lesion length (LL), Lesion area (LA) and Relative lesion area (RLA) growth (**A**), and chlorophyll content before Sclerotinia infection (SPADH), after infection (SPADI) and Chlorophyll Index (CI) values (**B**) in the most resistant and susceptible accessions of oil seed rape.
